# *de novo* assembly and population genomic survey of natural yeast isolates with the Oxford Nanopore MinION sequencer

**DOI:** 10.1101/066613

**Authors:** Benjamin Istace, Anne Friedrich, Léo d’Agata, Sébastien Faye, Emilie Payen, Odette Beluche, Claudia Caradec, Sabrina Davidas, Corinne Cruaud, Gianni Liti, Arnaud Lemainque, Stefan Engelen, Stefan Engelen, Patrick Wincker, Joseph Schacherer, Jean-Marc Aury

## Abstract

Oxford Nanopore Technologies Ltd (Oxford, UK) have recently commercialized MinION, a small and low-cost single-molecule nanopore sequencer, that offers the possibility of sequencing long DNA fragments. The Oxford Nanopore technology is truly disruptive and can sequence small genomes in a matter of seconds. It has the potential to revolutionize genomic applications due to its portability, low-cost, and ease of use compared with existing long reads sequencing technologies. The MinION sequencer enables the rapid sequencing of small eukaryotic genomes, such as the yeast genome. Combined with existing assembler algorithms, near complete genome assemblies can be generated and comprehensive population genomic analyses can be performed. Here, we resequenced the genome of the *Saccharomyces cerevisiae* S288C strain to evaluate the performance of nanopore-only assemblers. Then we *de novo* sequenced and assembled the genomes of 21 isolates representative of the *S. cerevisiae* genetic diversity using the MinION platform. The contiguity of our assemblies was 14 times higher than the Illumina-only assemblies and we obtained one or two long contigs for 65% of the chromosomes. This high continuity allowed us to accurately detect large structural variations across the 21 studied genomes. Moreover, because of the high completeness of the nanopore assemblies, we were able to produce a complete cartography of transposable elements insertions and inspect structural variants that are generally missed using a short-read sequencing strategy.

## Introduction

Today, long-read sequencing technology offers interesting alternatives to solve genome assembly difficulties and improve the completeness of genome assemblies, mostly in repetitive regions (Jain et al. 2015) where short-read sequencing has failed. Microbial or small eukaryotic genomes could now be fully assembled using Oxford Nanopore (Loman et al. 2015) or Pacific Biosciences reads alone (Chin et al. 2013; Koren and Phillippy 2015) or in combination with short but high quality reads (Koren et al. 2012; Goodwin et al. 2015; Madoui et al. 2015). Application of the single-molecule real-time (SMRT) sequencing platform to large complex eukaryotic genomes demonstrated the possibility of considerably improving genome assembly quality (Huddleston et al. 2014; Chaisson et al. 2015). Similar improvements were also accomplished using the 10x Genomics platform, and its application to the human genome produced encouraging results (Mostovoy et al. 2016; Zheng et al. 2016) and showed the importance of obtaining long and high-quality reads.

The most used sequencing technologies are based on the synthesis of new DNA strands, including the Illumina and Pacific Biosciences technologies (Mardis 2008). These sequencing technologies based on optical detection of nucleotide incorporations are often commercialized through large-sized and expensive instruments. For example, the cost of the commercially available Pacific Biosystems RS II instrument is high and the infrastructure and implementation needs make it inaccessible to large sections of the research community. This year Oxford Nanopore Technologies Ltd (ONT, Oxford, UK) commercialized MinION, a single-molecule nanopore sequencer that can be connected to a laptop through a USB interface (Loman and Watson 2015; Deamer et al. 2016). This system is portative (close to the size of a harmonica) and low-cost (currently USD 1,000 for the instrument). The MinION technology is based on an array of nanopores embedded on a chip that detects consecutive 6-mers of a single-strand DNA molecule by electrical sensing (Kasianowicz et al. 1996; Cherf et al. 2012; Manrao et al. 2012; Laszlo et al. 2014). In addition to its small size and low price, this new technology has several advantages over the older technologies. Library construction involves a simplified method, no amplification step is needed, and data acquisition and analyses occur in real time (Loose et al. 2016). Library preparation can be performed in two ways: (i) a 10-minute library preparation based on an enzymatic method for ‘1D’ sequencing (sequencing one strand of the DNA) or (ii) a library preparation based on ligation for ‘2D’ sequencing (sequencing both the template and complement strands of the DNA). In the 2D sequencing mode, the two strands of a DNA molecule are linked by a hairpin and sequenced consecutively. When the two strands of the molecule are read successfully, a consensus sequence is built to obtain a more accurate read (called 2D read). Otherwise only the template or complement strand sequence is provided (called 1D read).

Here, we sequenced the genomes of 22 *Saccharomyces cerevisiae* isolates to determine if the MinION system could be used in population genomic projects that require a deeper view of the genetic variation landscape. Even when the throughput of MinION was still heterogeneous, we were able to perform the sequencing in a reasonable time using six MinION devices. First, we resequenced the *Saccharomyces cerevisiae* S288C reference genome using a nanopore long-read sequencing strategy to evaluate recent assembly methods. We generated a complete benchmark of the assembly structures, as well as the completeness of complex regions. Next, we selected 21 strains of *S. cerevisiae* that were genetically diverse, based on preliminary results of the 1002 Yeast Genomes Project a large-scale short-read resequencing project (http://1002genomes.u-strasbg.fr/). The genomes of these 21 strains were *de novo* sequenced and assembled with Nanopore long-reads to have a better insight into the variation of their genomic architecture. We obtained near complete assembly, in terms of genes, as well as transposable elements and telomeric regions. The most contiguous assembly produced a single contig per chromosome, except for chromosomes 3 and 12, the latter contains the large repeated rDNA cluster.

## Results

### MinION data evaluation

We first sequenced the S288C genome by doing 11 MinION Mk1 runs with the R7.3 chemistry. On average, a 48-hours run produced more than 200 Mb of sequence, and the best run throughput was 400 Mb. Two 2D library types with 8 kb and 20 kb mean fragmentation sizes were used. They led to nearly 360,000 reads with a cumulative length of approximately 2.3 Gb and 63% of the nucleotides were in 2D reads, which represented a 187x and 118x genome coverage for 1D and 2D reads, respectively. Template reads had a median length of 8.9 kb while 2D reads had a median length of 7.7 kb. All sequencing reads were aligned to the S288C reference genome using BWA (Li and Durbin 2009) to assess their quality. We successfully aligned 95.6% of the 2D reads with an average error rate of 17.2% (Figure 1a). ONT tagged high-quality 2D reads as “2D pass” reads (reads with an average per-base quality higher than 9), and 99.7% of the 2D pass reads were aligned to the reference genome with an average error rate of 12.2%. We then parsed the alignment files to search for errors in stretches of the same nucleotide (homopolymers). About 85% of A, T, C, and G homopolymers of size 2 were present correctly in the reads. This percentage decreased rapidly to 65% for homopolymers of size 4 for A and T homopolymers and to 70% for C and G homopolymers. For size 7 homopolymers, it was 30% for A and T homopolymers and 35% for C and G homopolymers (**Figure S1a**).

**Figure 1:**
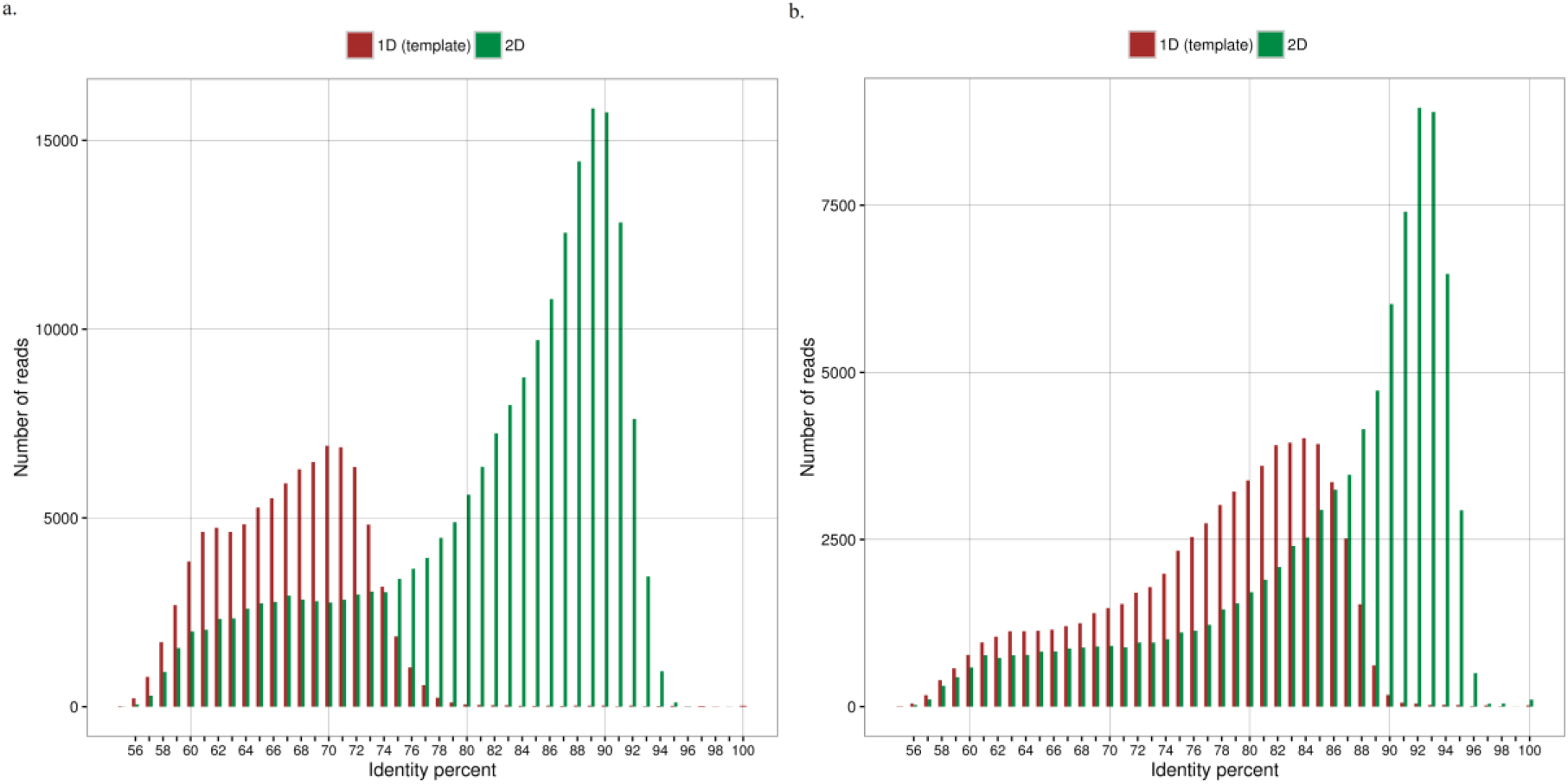
Identity distribution of Nanopore reads. Percent identity of the aligned MinION 1D (red bars) and 2D (green bars) reads. The MinION reads were aligned using LAST software. **a**. R7.3 chemistry **b**. R9 chemistry

We also sequenced the S288C genome using the R9 chemistry, the recently released version of the pore. We obtained approximately 1 Gb of reads; 568 Mb were 2D reads, which represents a 85x coverage with 1D reads and a 47x coverage with 2D reads. The mean 2D length was 6.1 kb. We aligned 82.1% of the 1D reads with a mean identity percentage of 82.8% and 94.3% of the 2D reads with a mean identity percentage of 85.2% (Figure 1b). As we did with the R7.3 reads, we also searched for errors in homopolymers (**Figure S1b**). The numbers of correct A, T, C, and G homopolymers started at about 90% for size equal to 2, then decreased to 75% for A and T homopolymers of size 4 and to 60% for the C and G homopolymers. For size 7 homopolymers, it was 32% for A, T, and C homopolymers and 35% for G homopolymers.

### Comparison of Nanopore-only assemblers

We tested Canu (Berlin et al. 2015), Miniasm (Li 2016), SMARTdenovo (Ruan) and ABruijn (Lin et al. 2016) with different subset of 1D, 2D, and 2D pass reads (**Supplementary File 2**) and kept the best assembly for each software.

With Canu, the best assembly was obtained with the whole set of 2D pass reads (67x coverage). The assembly was composed of 37 contigs with a cumulative length of 12 Mb and seven chromosomes were assembled in one or two contigs. After aligning the contigs to the S288C reference genome using Quast (Gurevich et al. 2013), we detected a high number of deletions (120,365), which were often localized in homopolymers (58%). As a consequence, only 454 of the 6,243 genes found in the assembly were insertion/deletion (indel)-free (**Table S1**). With Miniasm, the best assembly was obtained using the 2D reads corrected by Canu, which represented coverage of approximately 108x. The Miniasm assembly was composed of 28 contigs with a cumulative length of 11.8 Mb, and 13 chromosomes were assembled in one or two contigs. The consensus sequence contained a high proportion of mismatches and indels. With SMARTdenovo, 30x of the longest 2D reads produced the best assembly. It was composed of 26 contigs, with a total length of 12 Mb, and 14 chromosomes were assembled in one or two contigs. The SMARTdenovo assembly better covered the reference genome (>99%) and contained the highest number of genes (98.8% of the 6,350 S288C genes), but the Quast output again revealed a high number of deletions (128,050). With ABruijn, we obtained the best results using all the 2D reads as input, which represented coverage of approximately 120x. The assembly contained 23 contigs with a cumulative length of 11.9 Mb, and 14 chromosomes were assembled in one or two contigs (**Table S1**).

Next, we aligned the assemblies (Canu, Miniasm, SMARTdenovo, and ABruijn) to the S288C reference genome using NUCmer (Kurtz et al. 2004), and visualized the alignments with mummerplot (**Figures S2, S3, S4 and S5**). We also examined the coordinates of the alignments to search for chimera. We did not detect any chimeric contigs in the Canu, Miniasm, or SMARTdenovo assemblies; however, we did find some in the ABruijn assembly. Three chimeric contigs in the ABruijn assembly showed links between chromosomes 3 and 13 (first contig), chromosomes 3 and 2 (second contig), and chromosomes 10 and 2 (third contig). To verify that the portions of these contigs were effectively chimeric, we back aligned the Nanopore reads to the assembly and could not find any sequence that validated these links. Unsurprisingly, these three chimeric contigs were fused at Ty1 transposable element locations.

The alignment of each assembly to the reference genome showed that neither Canu, Miniasm, nor SMARTdenovo could assemble the mitochondrial (Mt) genome completely. Because ABruijn was the only assembler to assemble the complete Mt genome sequence, we decided to use it to assemble the Mt DNA of the remaining 21 yeast strains (see below).

Generally, long reads allow tandem duplicated genes to be resolve, as for instance the *CUP1* and *ENA1-2* gene families. We compared the maximum number of copies found in the Nanopore reads and the estimated number of copies based on Illumina reads coverage of these two tandem-repeated genes with the number of copies of these two genes in the four assemblies (**Table S2**). After aligning the paired-end reads to the reference sequence and computing of the coverage, we estimated that *CUP1* and *ENA1-2* were present in seven and four copies, respectively. The maximum numbers of copies of these genes in a single Nanopore read were eight for *CUP1* and five for *ENA1-2.* The numbers of copies of *CUP1* and *ENA1-2* were, respectively, nine and three in the Canu assembly, seven and two in the Miniasm assembly and seven and four in the SMARTdenovo and ABruijn assemblies.

The number of indels in each assembly was considerably high for each assembler. Thus, we tested Nanopolish (Loman et al. 2015), the most commonly used Nanopore-only error corrector. We used the SMARTdenovo assembly, which was the most continuous and gene-rich assembly and all 2D reads for this test. After the error correction step, the cumulative length of the contigs increased to 12.2 Mb and the N50 increased to 783 kb (at best it was 924 kb for the reference genome). The number of mismatches, insertions and deletions decreased to 1,930, 7,707, and 17,445 respectively. The number of genes increased to 6,273 complete and 2,590 without an indel (**Table S3**).

Although all metrics were improved, the number of indels still seemed too high, especially in the coding regions of the genes. We decided to polish all assemblies with 2x250bp Illumina paired-end reads, using Pilon (Walker et al. 2014), to verify if the general quality of the assembly improved. The polishing step increased the N50 of each assembly, and the maximum of 816 kb was obtained with the ABruijn assembly. Pilon reduced the number of errors of each assembly, and the Canu and ABruijn assemblies had the best base quality with about 16 mismatches (15.85 and 17.88 for Canu and ABruijn respectively) and 22 indels (22.49 and 21.76 for Canu and ABruijn respectively) per 100 kb. The SMARTdenovo assembly contained the highest number of complete genes (6,266) and the Canu assembly contained the highest number of genes without any indels (5,921) (Table 1).

**Table 1:** Metrics of the S288C assemblies after polishing. Assemblies were corrected using 300x of 2x250bp Illumina reads as input to Pilon. The resulting corrected assembly was then aligned to the S288C reference genome using Quast.

|  | Spades | Canu | Miniasm | SMARTdenovo | ABruijn |
| --- | --- | --- | --- | --- | --- |
| Reads dataset used | Illumina PE 2x250 bp | 2D pass | Canu-corrected | Longest 2D | 2D |
| Coverage | 300x | 67x | 108x | 30x | 120x |
| # reads > 10kb | 0 | 16,860 | 21,005 | 28,668 | 28,668 |
| # contigs | 376 | 37 | 28 | 26 | 23 |
| Cumulative size | 12,047,788 | 12,230,747 | 12,113,521 | 12,213,590 | 12,182,847 |
| Genome fraction (%) | 96.464 | 98.519 | 98.421 | 99.352 | 98.635 |
| N50 | 149,184 | 610,494 | 736,456 | 783,336 | 816,355 |
| N90 | 19,522 | 191,846 | 265,917 | 242,658 | 257,117 |
| L50 | 27 | 8 | 7 | 7 | 6 |
| L90 | 100 | 20 | 16 | 16 | 16 |
| # mismatches | 1,126 | 1,898 | 4,455 | 4,205 | 2,138 |
| # mismatches per 100 kb | 9.47 | 15.85 | 37.23 | 34.27 | 17.88 |
| # insertions | 81 | 1,657 | 3,164 | 2,384 | 1,325 |
| # deletions | 439 | 1,869 | 5,208 | 5,551 | 1,838 |
| # deletions in homopolymers | 38 | 868 | 4,248 | 4,023 | 740 |
| #indels per 100 kb | 1.97 | 22.49 | 57.27 | 46.76 | 21.76 |
| # genes | 6,087 + 177 partial | 6,241 + 32 partial | 6,215 + 37 partial | 6,266 + 33 partial | 6,243 + 45 partial |
| # genes without indels | 6,023 | 5,921 | 5,475 | 5,881 | 6,002 |

Finally, we evaluated the composition of each assembly for various elements (genes, repeated elements, centromeres and telomeric regions). We also generated an Illumina-only assembly using Spades assembler (Bankevich et al. 2012) to compare the number of features found in each assembly. All the assemblies contained nearly the same number of centromeres (120 bp regions in the reference genome assembly) and genes (Figure 2). The Nanopore assemblies contained between 45 and 50 Long Terminal Repeat (LTR) retrotransposons (average size of 5.8 kb), while the Illumina-only assembly contained only one. The smallest number of telomeres (three) was found in the ABruijn assembly, while nine, 18, 13, and 14 telomeres were found in the Illumina, Canu, Miniasm, and SMARTdenovo assemblies, respectively. The Illumina-only assembly contained five telomeric repeats (average size 100 bp), while the Nanopore-only assemblies contained between six and nine telomeric repeats. The ABruijn assembly contained the same number of genes encoded by the mitochondrial genome as the reference sequence because it was the only assembler to fully assemble the Mt genome.

**Figure 2:**
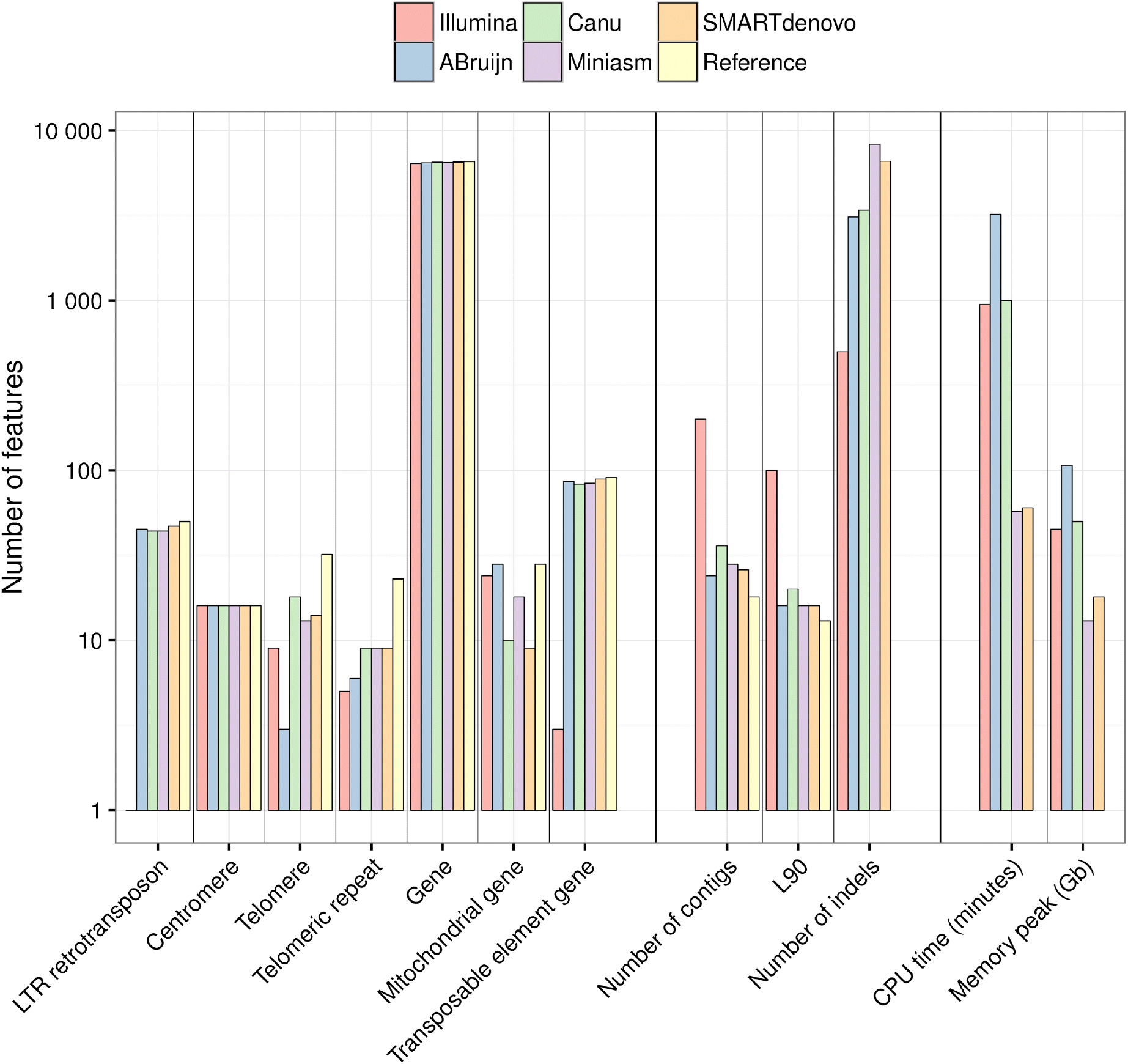
Feature composition of the S288C assemblies, continuity and quality metrics and assembler running statistics. The feature content of the best S288C assemblies for each assembler is shown in the left part of the figure. The feature composition was obtained by aligning each assembly to the S288C reference genome. Continuity and quality metrics for each assembly, obtained by using Quast, are shown in the middle part of the figure. The running time and the memory usage of each assembler are shown in the right part of the figure.

### S288C assemblies with R9 data

The R9 version of the pore was released too late for us to use it to sequence all the natural *S. cerevisiae* isolates. However, we did produce some data to compare the R7.3 and R9 assemblies. Because SMARTdenovo produced the best results, we used it to assemble the genome of the S288C strain. We input four different read datasets: all 1D and 2D reads, only 2D reads, 30x of the longest 2D reads or 30x of the longest 1D and 2D reads (**Table S4**).

This time, the 30x of the longest 1D and 2D reads dataset gave the best results. Indeed, the continuity of the assembly increased, and the number of contigs decreased from 26 with the R7.3 assembly to 23 with the R9 assembly. The number of indels also decreased from 133,676 with the R7.3 version to 95,012 with the R9 version. A direct consequence of using the R9 version was that almost all the genes were found, and 6,302 of the 6,350 known genes were complete and 1,226 did not contain any indels.

### Sequencing and assembly of the genomes of the 22 yeast strains

To explore the variability of the genomic architecture within *S. cerevisiae,* 21 natural isolates were sequenced in addition to the S288C reference genome using the same strategy, namely, a combination of long Nanopore and short Illumina reads. Sequenced isolates were selected to include as much diversity as possible in terms of global locations (including Europe, China, Brazil, and Japan), ecological sources (such as fermented beverages, dairy products, trees, fruit soil, and wine), as well as genetic variation highlighted in the frame of the extensive resequencing 1002 Yeast Genomes project (http://1002genomes.u-strasbg.fr/) (**Table S5**). Among these isolates, the nucleotide variability was distributed across 491,076 segregating sites and the genetic diversity, estimated by the average pairwise divergence (π), was 0.0062, which is close to what is observed for the whole species (Peter and Schacherer 2016).

A total of 78 MinION Mk1 runs were performed and the highest throughput we obtained was 650 Mb (1D and 2D reads). This led to 1.4 million of 2D reads with a cumulative length of 12 Gb. We obtained 2D coverage that ranged from 22x to 115x (**Figure S6**) among the strains with a median read length of approximately 5.4 kb and a maximum size of 75 kb (**Figure S7**). In general, three runs or less were sufficient to obtain the expected coverage. Next, for each strain, we gave varying coverages of the longest 2D reads as input to SMARTdenovo and retained the most contiguous assembly. These assemblies were then given as input to Pilon for a polishing step with around 300x of Illumina paired-end reads. After polishing, we obtained a median number of contigs of 27.5 (Table 2), the minimum number was for the CEI strain (18 contigs) and the maximum was for the BAM strain (105 contigs). The median cumulative length was 11.93 Mb and ranged from 11.83 Mb for the ADQ strain to 12.2 Mb for the CNT strain. The median N50 contig size was 593 kb and varied from 201 kb for the CIC strain to 896 kb for the ADQ strain. The L90 varied from 14 for the BCN, CEI, and CNT strains, to 72 for the BAM strain with a median equal to 19.5.

**Table 2:** Continuity metrics of the SMARTdenovo assemblies of all yeast strain genomes.

|  | # contigs | Cumul (bp) | N50 (bp) | N90 (bp) | L50 | L90 | Max size (bp) |
| --- | --- | --- | --- | --- | --- | --- | --- |
| ABH | 22 | 11,960,929 | 803,880 | 267,734 | 6 | 16 | 1,483,918 |
| ADM | 41 | 11,883,044 | 474,542 | 171,488 | 10 | 26 | 1,009,064 |
| ADQ | 26 | 11,828,347 | 896,166 | 223,992 | 6 | 18 | 1,223,692 |
| ADS | 33 | 11,706,636 | 524,733 | 247,699 | 9 | 21 | 1,050,223 |
| AEG | 23 | 12,026,175 | 681,360 | 273,814 | 7 | 16 | 1,244,014 |
| AKR | 25 | 11,911,766 | 729,090 | 243,900 | 7 | 17 | 1,056,085 |
| ANE | 47 | 11,900,397 | 312,705 | 144,286 | 11 | 31 | 933,716 |
| ASN | 40 | 11,904,493 | 394,798 | 143,405 | 11 | 28 | 846,371 |
| AVB | 31 | 11,991,127 | 609,633 | 199,011 | 7 | 20 | 1,225,549 |
| BAH | 28 | 11,829,394 | 571,862 | 227,561 | 8 | 20 | 1,066,359 |
| BAL | 27 | 11,907,375 | 678,155 | 269,114 | 7 | 19 | 1,075,839 |
| BAM | 105 | 11,996,380 | 162,412 | 53,623 | 24 | 72 | 450,388 |
| BCN | 19 | 11,775,292 | 785,507 | 458,793 | 6 | 14 | 1,410,650 |
| BDF | 45 | 12,068,568 | 460,458 | 116,953 | 10 | 29 | 863,099 |
| BHH | 26 | 11,973,506 | 577,727 | 221,661 | 7 | 18 | 1,530,377 |
| CBM | 68 | 11,553,446 | 258,798 | 86,167 | 16 | 44 | 521,412 |
| CEI | 18 | 11,987,201 | 800,227 | 451,575 | 6 | 14 | 1,480,681 |
| CFA | 24 | 11,834,226 | 726,317 | 225,716 | 7 | 17 | 1,032,352 |
| CFF | 81 | 12,162,869 | 236,957 | 83,285 | 18 | 54 | 550,022 |
| CIC | 96 | 12,016,445 | 201,870 | 63,799 | 22 | 63 | 377,026 |
| CNT | 22 | 12,171,929 | 800,046 | 440,742 | 6 | 14 | 1,402,970 |
| CRV (S288C) | 26 | 12,213,584 | 783,337 | 242,658 | 7 | 16 | 1,532,642 |
| Median | 27.5 | 11,936,347 | 593,680 | 224,854 | 7 | 19.5 | 1,061,222 |
| Reference | 17 | 12,157,105 | 924,431 | 439,888 | 6 | 13 | 1,531,933 |

To assemble the mitochondrial (Mt) genome, we used all the 2D reads as input to ABruijn. As a result, we obtained an assembly for each strain and extracted the Mt genome after mapping the contigs against the reference Mt genome. As was the case for the chromosomes, we used Pilon with Illumina paired-end reads to obtain a corrected consensus sequence.

### Transposable elements

The availability of high quality assemblies allowed us to establish an extensive map of the transposable elements (TEs) to obtain a global view of their content and positions within the 21 natural yeast isolates (Figure 3). Using a reference sequence for each of the five known TE families in yeast (namely Ty1 to Ty5), we mapped the TEs in each assembled genome. Among the 50 annotated TEs in the S288C reference genome, 47 were detected at the correct chromosomal locations in our assembly but three Ty1 locations were not recovered. Seven additional Ty1 elements were found at unannotated sites, three of them have already been detected in the reference genome (Bleykasten-Grosshans et al. 2011). These results attest to the high accuracy of our assembly strategy for TE detection and localization. Among the 22 isolates, the TE content was highly variable (Table 3), ranging from five to 55 elements, with a median value of 15. While the frequency of the Ty4 and Ty5 elements was clearly low in all the isolates (up to four and two elements, respectively), the Ty1, Ty2, and Ty3 elements were found in most of the isolates. The most abundant TEs were Ty1 and Ty2, except in the Chinese BAM isolate, in which 12 Ty3 elements were detected. As already described (Bleykasten-Grosshans et al. 2013), the pattern of insertion of these mobile elements is either specific to a given isolate, or shared by only a small number of isolates (mostly two or three). However, four insertion hotspots have been highlighted (shared by seven or more isolates) on chromosomes 2, 3, and 9. The shared insertion hotspots were generally not specific to a specific Ty family, except for the hotspot located on a subtelomeric region of the chromosome 3, which was specific to Ty5.

**Figure 3:**
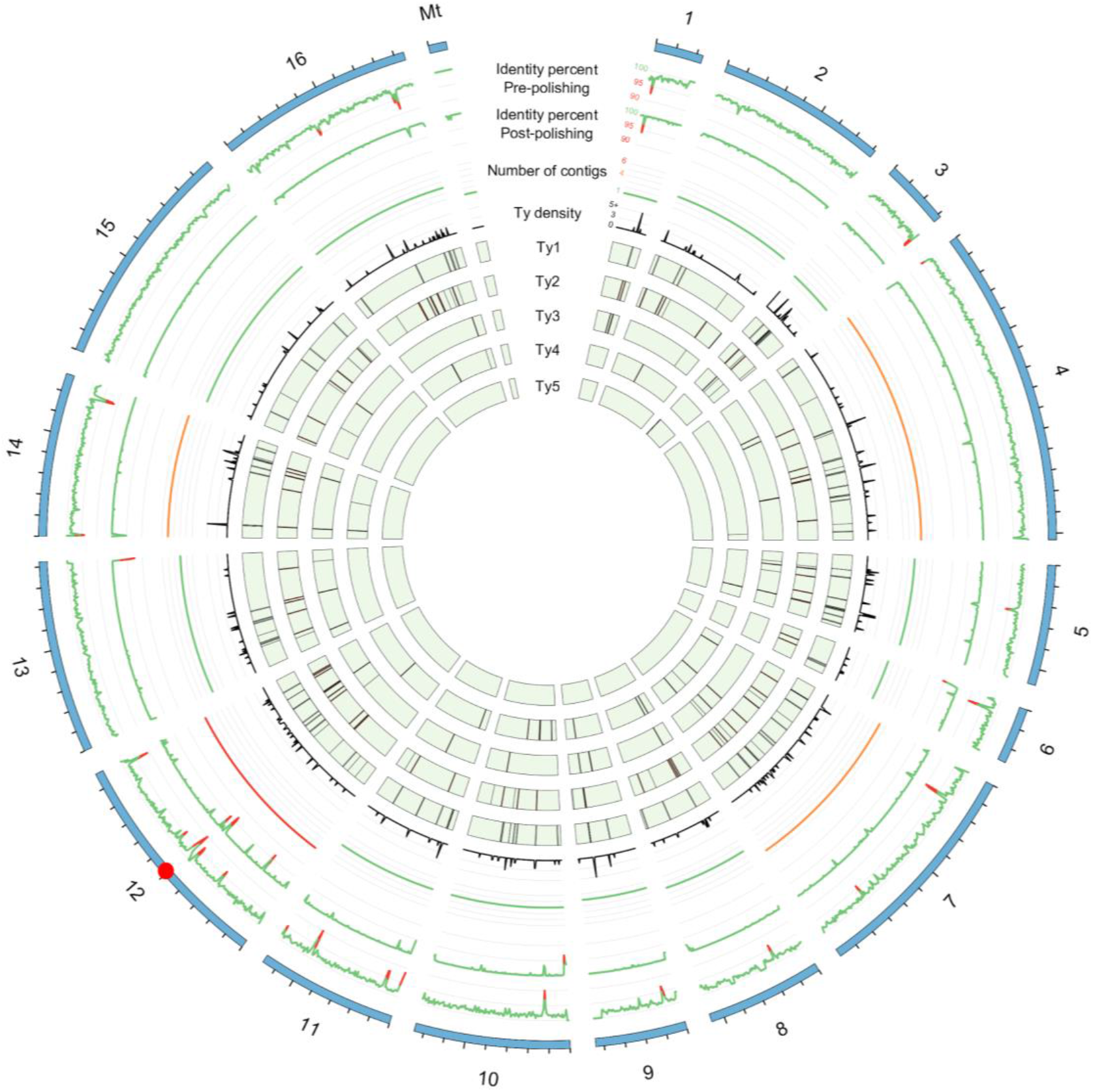
Cartography of the Ty transposon family. First and second tracks show, respectively, the percentage identity of the SMARTdenovo S288C assembly before and after polishing with Illumina paired-end reads using Pilon. The third track shows the 80^th^ percentile number of contigs obtained for each strain and for all chromosomes. The remaining tracks show the density of Ty transposons or positions of the Ty1, Ty2, Ty3, Ty4, and Ty5 transposons across all the yeast strains. The red dot on the karyotype track shows the position of the rDNA cluster.

**Table 3:** Number of copies of multiple transposons across all yeast strains assemblies.

|  | Ty1 | Ty2 | Ty3 | Ty4 | Ty5 |
| --- | --- | --- | --- | --- | --- |
| ABH | 4 | 7 | 6 | 3 | 2 |
| ADM | 5 | 8 | 1 | 1 | 0 |
| ADQ | 4 | 7 | 1 | 2 | 0 |
| ADS | 1 | 9 | 0 | 0 | 1 |
| AEG | 15 | 7 | 2 | 1 | 2 |
| AKR | 4 | 4 | 4 | 1 | 1 |
| ANE | 1 | 5 | 3 | 2 | 0 |
| ASN | 13 | 6 | 0 | 0 | 0 |
| AVB | 0 | 29 | 0 | 0 | 2 |
| BAH | 0 | 6 | 1 | 3 | 0 |
| BAL | 8 | 0 | 12 | 0 | 0 |
| BAM | 4 | 13 | 6 | 2 | 1 |
| BCN | 6 | 0 | 0 | 0 | 0 |
| BDF | 13 | 3 | 3 | 3 | 1 |
| BHH | 20 | 12 | 5 | 4 | 0 |
| CBM | 3 | 1 | 0 | 1 | 0 |
| CEI | 2 | 20 | 1 | 0 | 0 |
| CFA | 8 | 1 | 1 | 0 | 1 |
| CFF | 6 | 6 | 2 | 0 | 1 |
| CIC | 6 | 3 | 1 | 1 | 0 |
| CNT | 17 | 6 | 1 | 1 | 1 |
| CRV (S288C) | 36 | 13 | 2 | 3 | 1 |
| Reference | 31 | 13 | 2 | 3 | 1 |

### Structural variations

Structural variations such as copy number variants, large insertions and deletions, duplications, inversions and translocations are of great importance at the phenotypic variation level (Weischenfeldt et al. 2013). Compared with single nucleotide polymorphism (SNPs) and small indels, these variants are usually more difficult to identify, in particular because resequencing strategies have until recently focused mainly on the generation of short reads and reference-based genome analysis. Nanopore long reads sequencing data allow the copy numbers of tandem genes to be determined. As a testbed, we focused on two loci that are known to contain multi-copy genes, namely *ENA* and *CUP1. ENA* genes encode plasma membrane Na^+^-ATPase exporters, which play a role in the detoxification of Na+ ions in *S. cerevisiae. CUP1* genes encode metallothioneins, which bind copper and are involved in resistance to copper exposure by amplification of this locus. To determine the degree of divergence among the 21 strains, we searched for the numbers of copies of the *CUP1* and *ENA1-2* tandem-repeated genes in the assemblies (**Table S6**). For this purpose, we extracted the corresponding sequence from the S288C reference genome and aligned it to the assemblies of each strain. As expected and already reported (Strope et al. 2015), the copy numbers of *ENA1-2* and *CUP1* varied greatly across the strains. We found that the copy numbers of *ENA* genes in the 21 isolates ranged from 1 in 12 of the genomes to five in the BHH strain (**Table S6**). The copy numbers of *CUP1* genes fluctuated even more, ranging from one to 10 copies in the ABH and AEG strains. We also determined the fitness of the 21 isolates in the presence of CuSO_4_ and observed a correlation between the number of *CUP* genes and the resistance of the strain to high concentration of CuSO_4_ (Figure S8).

Besides copy number variants, we also focused on larger structural variants, such as translocations and inversions, because our highly contiguous assemblies allowed us to investigate these events. We aligned the polished assemblies of the 21 strains to the reference genome using NUCmer and inspected the alignments with the mummer software suite to search for structural variations. We detected 29 translocations and four inversions within the assemblies of 17 strains (**Table S7**). The median length of an inversion was 94 kb and their breakpoints were located mostly in intergenic regions. It is well recognized that SVs might play a major role in the genetic and phenotypic diversity in yeast (Hou et al. 2014; Naseeb et al. 2016). However, up to now, it was impossible to assemble and have an exhaustive view of the SVs content in any *S. cerevisiae* natural isolates. Indeed, short-read sequencing approaches are not suitable for SVs studies because they results in a high number of false positive as well as false negative detected events.

Among the detected events, one translocation detected between chromosomes 5 and 14 in the ABH isolate and another translocation between chromosomes 7 and 12 in the AVB isolate have already been described and confirmed in a reproductive isolation study in *S. cerevisiae* (Hou et al. 2014). A deeper investigation of our assemblies highlighted the presence of full-length Ty transposons at some junctions of the translocation events. For example, the complex Ty-rich junctions of the translocation between the chromosomes 7 and 12 in the ABH isolate was in complete accordance with previously reported results (Hou et al. 2014). Our results underline the high resolution of the constructed assemblies, and show that complex events, such as translocations, can be detected accurately with our strategy. Among the 22 isolates, six were devoid of translocation events whereas the other 16 carries one to four such structural rearrangements compared to the reference.

However, several limitations can be highlighted for these detections. Contrary to expectations, no translocation that specifically affected subtelomeric regions was identified, underlining the difficulty of discriminating regions that are variable and contain a large number of repeated segments. Moreover, the detection accuracy is highly dependent on the completeness of the assembly because, if translocation breakpoints are located on contigs boundaries, they will not be detectable.

### Mitochondrial genome variation

The ABruijn assembler allowed the construction of a single contig corresponding to the Mt genome for each isolate. To assess the quality of the assemblies, we aligned the polished S288C Mt contig to the reference sequence (GenBank: KP263414). Only four SNPs and a total of 15 bp long indels were detected. For all but two natural isolates, all the Mt genes (eight protein coding genes, two rRNA subunits and 24 tRNAs) were conserved and syntenous. The Mt genomes of the two remaining isolates (CNT and CFF) contained one and two repeated regions covering a total of 6.5 and 8 kb, respectively. In the CNT, the repeated region was in the *COX1* gene and affected its coding sequence. In the CFF isolate, the *COX1, ATP6,* and *ATP8* genes would have been tandemly duplicated. However, because we could not identify reads that clearly covered the repeated regions, we excluded these two Mt genome assemblies from our dataset.

The sizes of the 20 considered assemblies ranged from 73.5 to 86.9 kb, which is close to the size reported previously (Wolters et al. 2015). The differences in size between the assemblies can mainly be attributed to the intron content of the *COX1* and *COB* genes (from two to eight introns in *COX1* and from two to six introns in *COB).* These variations lead to extensive gene length variability ranging from 5.7 kb to 14.9 kb for *COX1* and from 3.2kb to 8.6 kb for *COB,* while the coding sequences of these 2 genes were exactly the same length among the 20 isolates. Intergenic regions also accumulate many small indels, including those that affect the interspersed GC-clusters, and a few large indels that sometimes correspond to variable hypothetical open reading frames (ORFs), leading to sizes that range from 51.6 to 58 kb. To a lesser extent, the 21S rRNA gene is also subjected to size variation that ranges from 3.2 to 4.4 kb.

## Discussion

One of the major advantages of the Oxford Nanopore technology is the possibility of sequencing very long DNA fragments. In our analyses, we obtained 2D reads up to 75 kb in length, indicating that the system was able to read without interruption a flow of at least 150,000 nucleotides. Furthermore, the results of this analysis indicate that the error rate of the ONT R7.3 reads was in the range that is obtained using existing long-read technologies (i.e, about 15% for 2D reads). However, the errors are not random and they significantly impact stretches of the same nucleotides (homopolymers), which seems to be a feature inherent to the ONT sequencing technology. Because the pore detects six nucleotides at a time, segmentation of events is problematic in genomic regions with homopolymers longer than six bases (David et al. 2016). With the current R7.3 release, homopolymers are prone to base deletion (representing 66% of the errors observed in homopolymers). It may be improved with a steadier passing speed through the pore or by increasing the speed of the molecule through the pore. In the same way, the basecaller algorithm could be optimized to increase the accuracy per base. ONT have recently reported several changes, including a fast mode (250 bp/second instead of 70 bp/second with R7.3 chemistry) and new basecaller software based on neural networks. These new features are incorporated in the R9 version of MinION. We performed R9 experiments, and observed a significant decrease in the error rate (with 1D and 2D reads, **Figure 1**). Using this new release, homopolymers were more prone to base insertions (representing 63% of the errors observed in homopolymers). Systematic errors are problematic for genome assembly because they lead to the construction of less accurate consensus sequences. Furthermore, indels negatively impact gene prediction because they can create frameshifts in the coding regions of genes. We concluded that nanopore-only assemblies are difficult to use for analysis at the gene level unless they are polished. However, polishing based only on nanopore reads was not sufficient because although it reduced the number of indels by more than seven times, we still had about 3,700 genes that were affected by potential frameshifts. The recently developed R9 chemistry greatly improved the overall quality of the consensus sequences, because starting with only 45x of 2D reads we obtained an assembly with the same contiguity but with a decrease of nearly 30% in the number of indels (95,012 compared with 133,676). We consider that the ONT sequencing platform will evolve in the coming years to produce high quality long reads. Until then, a mixed strategy using high quality short reads remains the only way to obtain high quality consensus sequences as well as a high level of contiguity. Indeed, for the assembly of repetitive regions, the nanopore-only assemblies outperformed the short-reads assemblies.

Our benchmark of nanopore-only assemblers shows that unfortunately a single “best assembler” does not exist. Canu reconstructed the telomeric regions better and provided a consensus of higher quality than Miniasm and SMARTdenovo. ABruijn seemed to produce the most continuous assembly but some of the contigs were chimeric. However, ABruijn was the only assembler to fully assemble the mitochondrial genome, and that is why we chose it to assemble the Mt genomes of the 22 yeast strains. SMARTdenovo provided good overall results for repetitive regions, completeness, contiguity, and speed. It was the most appropriate choice to assemble the genome of all the yeast strains even if its major drawback was the absence of the Mt genome sequence among the contig output.

The high contiguity of the 22 nanopore-only assemblies allowed us to detect transposable element insertions and to provide a complete cartography of these elements. Ty1 was the most abundant element and it was spread across the entire genome. Chromosome 12 was always the most fragmented in our assemblies due to the presence of the rDNA cluster (around 100 copies in tandem). Furthermore, we easily identified known translocations (between chromosomes 5 and 14 in the ABH isolate and between chromosomes 7 and 12 in the AVB isolate). The high contiguity of the assemblies seemed to be limited by the read size rather than the error rate. Work is still needed to prepare high-weight molecular DNA, enriched in long fragments. The yeast genomes were successfully assembled with 8 kb and 20kb fragment-sized libraries, but more complex genomes will require longer reads.

## Methods

### DNA extraction

Yeast cells were grown on YPD media (1% yeast extract, 2% peptone and 2% glucose) using liquid culture or solid plates. Total genomic DNA was purified from 30 ml YPD culture using Qiagen Genomic-Tips 100/G and Genomic DNA Buffers as per the manufacturer’s instructions. The quantity and quality of the extracted DNA were controlled by migration on agarose gel, spectrophotometry (NanoDrop ND-1000), and fluorometric quantification (Qubit, ThermoFisher).

### Illumina PCR-free library preparation and sequencing

DNA (6 μg) was sonicated to a 100 to 1500 bp size range using a Covaris E210 sonicator (Covaris, Woburn, MA, USA). Fragments were end-repaired using the NEBNext^®^ End Repair Module (New England Biolabs, Ipswich, MA, USA) and 3'-adenylated with the NEBNext^®^ dA-Tailing Module. Illumina adapters were added using the NEBNext® Quick Ligation Module. Ligation products were purified with AMPure XP beads (Beckmann Coulter Genomics, Danvers, MA, USA). Libraries were quantified by qPCR using the KAPA Library Quantification Kit for Illumina Libraries (KapaBiosystems, Wilmington, MA, USA) and library profiles were assessed using a DNA High Sensitivity LabChip kit on an Agilent Bioanalyzer (Agilent Technologies, Santa Clara, CA, USA). Libraries were sequenced on an Illumina MiSeq or a HiSeq 2500 instrument (San Diego, CA, USA) using 300 or 250 base-length read chemistry in a paired-end mode.

### Nanopore 20 kb libraries preparation

MinION sequencing libraries were prepared according to the SQK-MAP005 or SQK-MAP006-MinION gDNA Sequencing Kit protocols. Six to 10 μg of genomic DNA was sheared to approximately 20,000 bp with g-TUBE (Covaris, Woburn, MA, USA). After clean-up using 0.4x AMPure XP beads, sequencing libraries were prepared according to the SQK-MAP005 or SQK-MAP006 Sequencing Kit protocols, including the PreCR treatment (NEB, Ipswich, USA) for the SQK-MAP005 protocol or the NEBNext FFPE DNA repair step (NEB, Ipswich, USA) for the SQK-MAP006 protocol.

### Nanopore 8 kb libraries preparation

MinION sequencing libraries were prepared according to the SQK-MAP005 or SQK-MAP006-MinION gDNA Sequencing Kit protocols. Two μg of genomic DNA was sheared to approximately 8,000 bp with g-TUBE. After clean-up using 1x AMPure XP beads, sequencing libraries were prepared according to the SQK-MAP005 or SQK-MAP006 Sequencing Kit protocol, including the PreCR treatment for the SQK-MAP005 protocol or the NEBNext FFPE DNA repair step for the SQK-MAP006 protocol.

### Nanopore Low input 8 kb libraries preparation

The following protocol was applied to some samples. Five hundred ng of genomic DNA was sheared to approximately 8,000 bp with g-TUBE. After clean-up using 1x AMPure XP beads and the NEBNext FFPE DNA repair step, 100 ng of DNA was prepared according to the Low Input Expansion Pack Protocol for genomic DNA.

### MinION^TM^ flow cell preparation and sample loading

The sequencing mix was prepared with 8 μL of the DNA library, water, the Fuel Mix and the Running buffer according to the SQK-MAP005 or the SQK-MAP006 protocols. The sequencing mix was added to the R7.3 flowcell for a 48 hours run. The flowcell was then reloaded three times according to the following schedule: 5 hours (4 μL of DNA library), 24 hours (8 μL of DNA library) and 29 hours (4 μL of DNA library). Regarding the Low Input libraries, the flowcell was loaded and then reloaded after 24 hours of run time with a sequencing mix containing 10 μL of the DNA library.

### MinION^®^ sequencing and reads filtering

Read event data generated by MinKNOW^TM^ control software (version 0.50.1.15 to 0.51.1.62) were base-called using the Metrichor^TM^ software (version 2.26.1 to 2.38.3). The data generated (pores metrics, sequencing, and base-calling data) by MinION software were stored and organized using a Hierarchical Data Format (HDF5). Three types of reads were obtained: template, complement, and two-directions (2D). The template and complement reads correspond to sequencing of the two DNA strands. Metrichor^TM^ combines template and complement reads to produce a consensus (2D) sequence (Quick et al. 2014). FASTA reads were extracted from MinION HDF5 files using poretools (Loman and Quinlan 2014). To assess the quality of the MinION reads, we aligned reads against the *S. cerevisiae* S288C reference genome using the LAST aligner (version 588) (Kielbasa et al. 2011). Because the MinION reads are long and have a high error rate we used a gap open penalty of 1 and a gap extension of 1.

### Illumina reads processing and quality filtering

After the Illumina sequencing, an in-house quality control process was applied to the reads that passed the Illumina quality filters. The first step discards low-quality nucleotides (Q<20) from both ends of the reads. Next, Illumina sequencing adapters and primers sequences were removed from the reads. Then, reads shorter than 30 nucleotides after trimming were discarded. These trimming and removal steps were achieved using in-house-designed software based on the FastX package (FASTX-Toolkit). The last step identifies and discards read pairs that mapped to the phage phiX genome, using SOAP (Li et al. 2009b) and the phiX reference sequence (GenBank: NC_001422.1). This processing resulted in high-quality data and improvement of the subsequent analyses.

### Assembler evaluation

To determine the assembler to use on the *de novo* sequenced 22 yeast strains, tests were conducted on S288C, the only *S. cerevisiae* strain for which there is an established reference genome. We used different subsets of the reads as input to Canu (github commit ae9eecc), Miniasm (github commit 17d5bd1), SMARTdenovo (github commit 61cf13d), and ABruijn (github commit dc209ee), four assemblers that can take advantage of long reads. These subsets consisted of varying coverages of 1D, 2D, 2D pass reads, which are 2D reads that have an average quality greater than nine, and reads corrected by Canu. Canu was executed with the following parameters: genomeSize=12m, minReadLength=5000, mhapSensitivity=high, corMhapSensitivity=high, errorRate=0.01 and corOutCoverage=500. Miniasm was run with the default parameters indicated on the github website. SMARTdenovo was executed with the default parameters and -c 1 to run the consensus step. ABruijn was run with default parameters. After the assembly step, we polished each set of contigs with Pilon, using 300X of Illumina 2x250 bp paired-end reads. Assemblies were aligned to the S288C reference genome using Quast in conjunction with the GFF file of S288C to detect assembly errors, and complete and partial genes. We also visualized the alignments using mummerplot to detect chimeric contigs.

### Genes and transposons detection

To detect genes and transposons in the assemblies, we extracted the corresponding sequences from the reference genome. We then mapped these elements to the assemblies using the Last aligner. Only alignments that showed more than 80% identity over at least 90% of the sequence length were retained and considered as a match. We used a similar procedure to count the maximum number of gene in the Nanopore reads dataset, the only modification was that the percentage identity had to be at least 70% to account for the high error rate of the reads. To estimate the number of copies in the Illumina reads, we aligned paired-end reads to the reference genome with BWA aln and then computed the coverage using samtools mpileup algorithm (Li et al. 2009a) and divided the number we obtained for each region of interest by the median coverage of the corresponding chromosome.

### Feature number estimation

We generated an Illumina-only assembly using Spades version v3.7.0 with default parameters and compare the completeness of this assembly to the nanopore-only assemblies. To estimate the number of features across all S288C assemblies, we aligned each post-polishing consensus sequence to the S288C reference genome using NUCmer. Only the best alignments were conserved by using the *delta-filter −1* command. Next, we used the bedtools suite (Quinlan and Hall 2010) with the command *bedtools intersect -u -wa -f 0.99* to compare the alignments to the reference GFF file. Finally, we counted the number of features of our interest.

### Circularization of mitochondrial genomes

To circularize the Mt genomes, we split the contig corresponding to the Mt sequence in each strain into two distinct contigs. Then, we gave the two contigs as input to the minimus2 (Schatz et al. 2013) tool from the AMOS package. As a result, we obtained a single contig that did not contain the overlap corresponding to the circularization zone. Finally, to start the Mt sequence of all isolates at the same position as the reference, we mapped each Mt sequence to the reference using NUCmer. The *show-coords* command allowed us to identify the position in the Mt sequences of all the strains that corresponded to the first position of the reference Mt genome.

## Data accessibility

The 22 genome assemblies are freely available at http://www.genoscope.cns.fr/yeast. The Illumina and MinION data are available in the European Nucleotide Archive under accession number ERP016443.

## Additional files

All the supporting data are included as a two additional files: a first one which contains Figures S1-S8 and Tables S1-S7 and an excel file which contains the metrics of all assemblies generated in this study.

## Abbreviations

ONT: Oxford Nanopore Technology
SMRT: Single-Molecule Real-Time Sequencing
USB: Universal Serial Bus
Mt: Mitochondrial
LTR: Long Terminal Repeat
SNP: Single Nucleotide Polymorphism
ORF: Open Reading Frame
MAP: MinION Access Programme

## Author’s contributions

CCA extracted the DNA. EP, OB, CCR and AL optimized and performed the sequencing. BI, AF, LDA, SF, SD, SE and JMA performed the bioinformatic analyses. BI, AF, JS and JMA wrote the article. GL, PW, JS and JMA supervised the study.

## Acknowledgements

This work was supported by the Genoscope, the Commissariat à l’Energie Atomique et aux Energies Alternatives (CEA), and France Génomique (ANR-10-INBS-09-08). The authors are grateful to Oxford Nanopore Technologies Ltd for providing early access to the MinION device through the MinION Access Programme (MAP) and we thank the staff of Oxford Nanopore Technology Ltd for technical help. The authors acknowledge Pierre Le Ber and Claude Scarpelli for continuous support.

## References

Bankevich A, Nurk S, Antipov D, Gurevich AA, Dvorkin M, Kulikov AS, Lesin VM, Nikolenko SI, Pham S, Prjibelski AD et al. 2012. SPAdes: a new genome assembly algorithm and its applications to single-cell sequencing. Journal of computational biology : a journal of computational molecular cell biology 19(5): 455–477.

Berlin K, Koren S, Chin CS, Drake JP, Landolin JM, Phillippy AM. 2015. Assembling large genomes with single-molecule sequencing and locality-sensitive hashing. Nature biotechnology 33(6): 623–630.

Bleykasten-Grosshans C, Friedrich A, Schacherer J. 2013. Genome-wide analysis of intraspecific transposon diversity in yeast. BMC genomics 14: 399.

Bleykasten-Grosshans C, Jung PP, Fritsch ES, Potier S, de Montigny J, Souciet JL. 2011. The Ty1 LTR-retrotransposon population in Saccharomyces cerevisiae genome: dynamics and sequence variations during mobility. FEMS yeast research 11(4): 334–344.

Chaisson MJ, Huddleston J, Dennis MY, Sudmant PH, Malig M, Hormozdiari F, Antonacci F, Surti U, Sandstrom R, Boitano M et al. 2015. Resolving the complexity of the human genome using single-molecule sequencing. Nature 517(7536): 608–611.

Cherf GM, Lieberman KR, Rashid H, Lam CE, Karplus K, Akeson M. 2012. Automated forward and reverse ratcheting of DNA in a nanopore at 5-A precision. Nature biotechnology 30(4): 344–348.

Chin CS, Alexander DH, Marks P, Klammer AA, Drake J, Heiner C, Clum A, Copeland A, Huddleston J, Eichler EE et al. 2013. Nonhybrid, finished microbial genome assemblies from long-read SMRT sequencing data. Nature methods 10(6): 563–569.

David M, Dursi LJ, Yao D, Boutros PC, Simpson JT. 2016. Nanocall: An Open Source Basecaller for Oxford Nanopore Sequencing Data. bioRxiv.

Deamer D, Akeson M, Branton D. 2016. Three decades of nanopore sequencing. Nature biotechnology 34(5): 518–524.

FASTX-Toolkit. http://hannonlab.cshl.edu/fastxtoolkit/.

Goodwin S, Gurtowski J, Ethe-Sayers S, Deshpande P, Schatz MC, McCombie WR. 2015. Oxford Nanopore sequencing, hybrid error correction, and de novo assembly of a eukaryotic genome. Genome research 25(11): 1750–1756.

Gurevich A, Saveliev V, Vyahhi N, Tesler G. 2013. QUAST: quality assessment tool for genome assemblies. Bioinformatics 29(8): 1072–1075.

Hou J, Friedrich A, de Montigny J, Schacherer J. 2014. Chromosomal rearrangements as a major mechanism in the onset of reproductive isolation in Saccharomyces cerevisiae. Current biology : CB 24(10): 1153–1159.

Huddleston J, Ranade S, Malig M, Antonacci F, Chaisson M, Hon L, Sudmant PH, Graves TA, Alkan C, Dennis MY et al. 2014. Reconstructing complex regions of genomes using long-read sequencing technology. Genome research 24(4): 688–696.

Jain M, Fiddes IT, Miga KH, Olsen HE, Paten B, Akeson M. 2015. Improved data analysis for the MinION nanopore sequencer. Nature methods 12(4): 351–356.

Kasianowicz JJ, Brandin E, Branton D, Deamer DW. 1996. Characterization of individual polynucleotide molecules using a membrane channel. Proceedings of the National Academy of Sciences of the United States of America 93(24): 13770–13773.

Kielbasa SM, Wan R, Sato K, Horton P, Frith MC. 2011. Adaptive seeds tame genomic sequence comparison. Genome research 21(3): 487–493.

Koren S, Phillippy AM. 2015. One chromosome, one contig: complete microbial genomes from long-read sequencing and assembly. Current opinion in microbiology 23: 110–120.

Koren S, Schatz MC, Walenz BP, Martin J, Howard JT, Ganapathy G, Wang Z, Rasko DA, McCombie WR, Jarvis ED et al. 2012. Hybrid error correction and de novo assembly of single-molecule sequencing reads. Nature biotechnology 30(7): 693–700.

Kurtz S, Phillippy A, Delcher AL, Smoot M, Shumway M, Antonescu C, Salzberg SL. 2004. Versatile and open software for comparing large genomes. Genome biology 5(2): R12.

Laszlo AH, Derrington IM, Ross BC, Brinkerhoff H, Adey A, Nova IC, Craig JM, Langford KW, Samson JM, Daza R et al. 2014. Decoding long nanopore sequencing reads of natural DNA. Nature biotechnology 32(8): 829–833.

Li H. 2016. Minimap and miniasm: fast mapping and de novo assembly for noisy long sequences. Bioinformatics 32(14): 2103–2110.

Li H, Durbin R. 2009. Fast and accurate short read alignment with Burrows-Wheeler transform. Bioinformatics 25(14): 1754–1760.

Li H, Handsaker B, Wysoker A, Fennell T, Ruan J, Homer N, Marth G, Abecasis G, Durbin R. 2009a. The Sequence Alignment/Map format and SAMtools. Bioinformatics 25(16): 2078–2079.

Li R, Yu C, Li Y, Lam TW, Yiu SM, Kristiansen K, Wang J. 2009b. SOAP2: an improved ultrafast tool for short read alignment. Bioinformatics 25(15): 1966–1967.

Lin Y, Yuan J, Kolmogorov M, Shen MW, Pevzner PA. 2016. Assembly of Long Error-Prone Reads Using de Bruijn Graphs. bioRxiv.

Loman NJ, Quick J, Simpson JT. 2015. A complete bacterial genome assembled de novo using only nanopore sequencing data. Nature methods 12(8): 733–735.

Loman NJ, Quinlan AR. 2014. Poretools: a toolkit for analyzing nanopore sequence data. Bioinformatics 30(23): 3399–3401.

Loman NJ, Watson M. 2015. Successful test launch for nanopore sequencing. Nature methods 12(4): 303–304.

Loose M, Malla S, Stout M. 2016. Real-time selective sequencing using nanopore technology. Nature methods.

Madoui MA, Engelen S, Cruaud C, Belser C, Bertrand L, Alberti A, Lemainque A, Wincker P, Aury JM. 2015. Genome assembly using Nanopore-guided long and error-free DNA reads. BMC genomics 16: 327.

Manrao EA, Derrington IM, Laszlo AH, Langford KW, Hopper MK, Gillgren N, Pavlenok M, Niederweis M, Gundlach JH. 2012. Reading DNA at single-nucleotide resolution with a mutant MspA nanopore and phi29 DNA polymerase. Nature biotechnology 30(4): 349–353.

Mardis ER. 2008. Next-generation DNA sequencing methods. Annual review of genomics and human genetics 9: 387–402.

Mostovoy Y, Levy-Sakin M, Lam J, Lam ET, Hastie AR, Marks P, Lee J, Chu C, Lin C, Dzakula Z et al. 2016. A hybrid approach for de novo human genome sequence assembly and phasing. Nature methods 13(7): 587–590.

Naseeb S, Carter Z, Minnis D, Donaldson I, Zeef L, Delneri D. 2016. Widespread Impact of Chromosomal Inversions on Gene Expression Uncovers Robustness via Phenotypic Buffering. Molecular biology and evolution 33(7): 1679–1696.

Peter J, Schacherer J. 2016. Population genomics of yeasts: towards a comprehensive view across a broad evolutionary scale. Yeast 33(3): 73–81.

Quick J, Quinlan AR, Loman NJ. 2014. A reference bacterial genome dataset generated on the MinION portable single-molecule nanopore sequencer. GigaScience 3: 22.

Quinlan AR, Hall IM. 2010. BEDTools: a flexible suite of utilities for comparing genomic features. Bioinformatics 26(6): 841–842.

Ruan J. https://github.com/ruanjue/smartdenovo.

Schatz MC, Phillippy AM, Sommer DD, Delcher AL, Puiu D, Narzisi G, Salzberg SL, Pop M. 2013. Hawkeye and AMOS: visualizing and assessing the quality of genome assemblies. Briefings in bioinformatics 14(2): 213–224.

Strope PK, Skelly DA, Kozmin SG, Mahadevan G, Stone EA, Magwene PM, Dietrich FS, McCusker JH. 2015. The 100-genomes strains, an S. cerevisiae resource that illuminates its natural phenotypic and genotypic variation and emergence as an opportunistic pathogen. Genome research 25(5): 762–774.

Walker BJ, Abeel T, Shea T, Priest M, Abouelliel A, Sakthikumar S, Cuomo CA, Zeng Q, Wortman J, Young SK et al. 2014. Pilon: an integrated tool for comprehensive microbial variant detection and genome assembly improvement. PloS one 9(11): e112963.

Weischenfeldt J, Symmons O, Spitz F, Korbel JO. 2013. Phenotypic impact of genomic structural variation: insights from and for human disease. Nature reviews Genetics 14(2): 125–138.

Wolters JF, Chiu K, Fiumera HL. 2015. Population structure of mitochondrial genomes in Saccharomyces cerevisiae. BMC genomics 16: 451.

Zheng GX, Lau BT, Schnall-Levin M, Jarosz M, Bell JM, Hindson CM, Kyriazopoulou-Panagiotopoulou S, Masquelier DA, Merrill L, Terry JM et al. 2016. Haplotyping germline and cancer genomes with high-throughput linked-read sequencing. Nature biotechnology 34(3): 303–311.

